# Perimenopause promotes neuroinflammation in select hippocampal regions in a mouse model of Alzheimer’s disease

**DOI:** 10.1101/2025.03.14.643317

**Authors:** Roberta Marongiu, Jimcy Platholi, Laibak Park, Fangmin Yu, Garrett Sommer, Clara Woods, Teresa A. Milner, Michael J. Glass

**Author notes:** Corresponding author for manuscript: Dr. Teresa A. Milner, Feil Family Brain and Mind Research Institute Weill Cornell Medicine, 407 East 61st Street, RM 307 New York, NY 10065 USA. Corresponding authors, communications: Teresa A. Milner - Michael J. Glass. These authors contributed equally to this work and share first authorship (Roberta Marongiu, Jimcy Platholi). These authors contributed equally to this work and share senior authorship (Teresa A. Milner, Michael J. Glass). **Submission for 2025 Special Issue:** “Estrogens and Neurodegeneration: a Link Between Menopause and Alzheimer’s Diseases in Women” hosted by Frontiers in Molecular Biosciences. **Conflict of interest:** The authors declare no competing financial interests.

## Abstract

Alzheimer’s disease (AD) is the most common neurodegenerative disorder characterized by age-dependent amyloid beta (Aβ) aggregation and accumulation, neuroinflammation, and cognitive deficits. Significantly, there are prominent sex differences in the risk, onset, progression, and severity of AD, as well as response to therapies, with disease burden disproportionately affecting women. Although menopause onset (i.e., perimenopause) may be a critical transition stage for AD susceptibility in women, the role of early ovarian decline in initial disease pathology, particularly key neuroinflammatory processes, is not well understood. To study this, we developed a unique mouse model of perimenopausal AD by combining an accelerated ovarian failure (AOF) model of menopause induced by 4-vinylcyclohexene diepoxide (VCD) with the 5xFAD transgenic AD mouse model. To target early stages of disease progression, 5xFAD females were studied at a young age (∼4 months) and at the beginning stage of ovarian failure analogous to human perimenopause (termed “peri-AOF”), and compared to age-matched males. Assessment of neuropathology was performed by immunohistochemical labeling of Aβ as well as markers of astrocyte and microglia activity in the hippocampus, a brain region involved in learning and memory that is deleteriously impacted during AD. Our results show that genotype, AOF, and sex contributed to AD-like pathology. Aggregation of Aβ was heightened in female 5xFAD mice and further increased at peri-AOF, with hippocampal subregion specificity. Further, select increases in glial activation also paralleled Aβ pathology in distinct hippocampal subregions. However, cognitive function was not affected by peri-AOF. These findings align with the hypothesis that perimenopause constitutes a period of susceptibility for AD pathogenesis in women.

## INTRODUCTION

Dementia is a leading contributor to the global burden of disease [1] with Alzheimer’s disease (AD) constituting approximately 50-70% of cases [2]. AD is characterized by progressive neurodegeneration and cognitive dysfunction [3, 4]. A main hallmark of AD neuropathology is the accumulation of parenchymal plaques containing aggregated amyloid-beta (Aβ) in the cerebral cortex and the hippocampal formation [5]. In addition to Aβ deposition, AD involves a complex set of related neurodegenerative processes including neuroinflammation [6]. In AD animal models, neuroinflammation characterized by the activation of reactive astrocytes and microglia [7] is one of the earliest pathological manifestations, likely contributing to synaptic and neuronal loss.

Important sex differences in AD are well documented, with women experiencing a disproportionately greater disease burden [8]. The incidence of AD is at least two-fold higher in women compared to men [2] with women exhibiting faster disease progression [9] and greater cognitive impairment at comparable stages of AD [10]. Perimenopause, the transitional phase of irregular gonadal hormonal production and cycling before full menopause, may be a particularly vulnerable period for the onset of mild cognitive impairment and AD. This is supported by evidence that low endogenous estrogen levels are associated with increased AD risk [11], and that early or late age at menopause is associated with an elevated or decreased risk for AD, respectively [12–14]. Furthermore, there is growing evidence that initiating estrogen replacement soon after menopause may help mitigate dementia development [15]. Significantly, perimenopause is also associated with declines in brain volume [16] and increases in Aβ expression [16, 17]. Despite these associations, the mechanisms underlying the heightened perimenopausal risk for AD, particularly those related to neuroinflammation and cognitive decline in the hippocampus, remain unclear.

Animal models may help elucidate the mechanisms driving perimenopausal AD risk. The commonly used 5xFAD transgenic mouse model, which expresses five familial mutations in two AD risk genes, exhibits increased Aβ production and plaque formation that parallels AD pathology [18]. Notably, sex differences have been reported in 5xFAD mice, with females showing earlier increases in inflammatory gene expression [19], glial markers [20, 21] as well as Aβ levels [19, 20, 22]. This neuropathology correlates with worse cognitive performance, in some studies [19–21], but not all [23, 24]. Markers of brain inflammation appear as early as three months of age, suggesting that sex differences in AD-like pathology emerge at a prodromal stage [19]. Additionally, female 5xFAD mice show increased hippocampal Aβ and elevated expression of immune-related genes and proteins compared to age-matched males [18, 25–29]. These sex differences may be linked to changes in estrogen signaling [30–32]. Thus, combining mouse models of AD with perimenopause in females may help to isolate the effects of hormonal changes on AD neuropathology.

The 4-vinylcyclohexene diepoxide (VCD) ovatoxin mimics perimenopause in rodents by producing accelerated ovarian failure (AOF) paralleling the irregular hormone fluctuations seen during human perimenopause (termed “peri-AOF” in rodents) before transitioning to full menopause [33–35]. This model allows for controlled induction of AOF at various times/ages following sexual maturity in younger animals, reducing the confounding effects of chronological aging, a main AD risk factor [33–35]. The VCD model has been used to investigate how ovarian failure influences metabolic [36–38], aging [39], and cerebrovascular [40] factors to influence cognitive function, brain plasticity and amyloid pathology in wild-type (WT) mice or in mouse models of cognitive impairment and dementia. However, the impact of early ovarian failure (peri-AOF) on Aβ-related pathology and neuroinflammatory response in 5xFAD mice is unknown.

We investigated whether peri-AOF contributes to hippocampal neuroinflammation in young 5xFAD mice treated with VCD. A granular assessment of Aβ levels and glial markers of neuroinflammation was conducted by analyzing all major hippocampal subfields at rostral and caudal levels. Cognitive performance was assessed using tests of learning and memory. To characterize early AD-like neuropathology, we focused on young (∼4 months) 5xFAD mice. Age-matched male mice were also tested to evaluate the effect of biological sex.

## EXPERIMENTAL PROCEDURES

### ANIMALS

Young adult (∼2 month-old at the initiation of the experiments [41]) C57BL/6 WT mice (n= 22 females and n= 11males) and transgenic 5xFAD (C57BL/6 background; n= 22 female and n= 11 male) mice were bred and maintained in a colony at Weill Cornell Medicine (WCM). Breeding pairs of hemizygous 5xFAD mice were obtained from the Jackson Laboratory (Bar Harbor, ME; JAX MMRRC stock#034840). Mice were bred in-house and genotyped prior to experimentation. 5xFAD mice express human APP and PSEN1 transgenes with a total of five AD-linked mutations under the control of the Thy1 promoter. This mouse model exhibits early-onset parenchymal Aβ aggregation correlated with cognitive deficits [42, 43]. Amyloid deposition begins in the cerebral parenchyma at 2-3 months of age, with little accumulation in the cerebral vasculature, and amyloid plaques are found throughout the hippocampus and cortex by six months [18, 44] . Astrogliosis and microgliosis begin around two months, developing in parallel with plaque deposition [18]. Mice were housed in groups of three to four animals per cage and maintained on a 12-hr light/dark cycle (lights out 18:00 hours) with *ad libitum* access to water and rodent chow. At euthanasia, mice weighed 23-32 grams. All experiments were approved by the WCM Institutional Animal Care and Use Committees and followed the National Institutes of Health guidelines for the Care and Use of Laboratory Animals guidelines.

#### AOF model of perimenopause

AOF induction by VCD treatment in mice has been shown to recapitulate the gradual hormonal fluctuations that correspond to peri- and post-menopause in humans (reviewed in [45, 46]). The AOF model can be used to separate hormonal effects from aging effects and can be applied to any mouse genotype [33–35]. Low dose VCD injections selectively deplete ovarian primary follicles without negatively affecting peripheral tissues, kidney, or liver function [34, 47–49]. VCD does not directly increase inflammation markers in the brain, including the hippocampus [33].

##### AOF induction

Gonadally intact 53-58-postnatal-day-old female mice received 130 mg/kg VCD (cat. # S453005 Millipore Sigma, St. Louis, MO) in sesame oil (cat. # 8008-74-0 Millipore Sigma) for 5 days per week for 3 weeks [35, 45]. Control mice received injections of sesame oil only. Prior studies from our lab and others [33, 47, 50] established the peri-AOF stage as occurring 58 days after the first VCD injection. At this stage (∼3.5 months old), the mice exhibit irregular, prolonged estrous cycles and elevated plasma follicle stimulating hormone [50–52]. Behavioral assessments of VCD- and oil-treated females, as well as aged-matched males, were initiated when mice were about 4 months of age (the peri-AOF stage of VCD mice). A timeline of the experimental procedures is shown in Figure 1.

**Figure 1:**
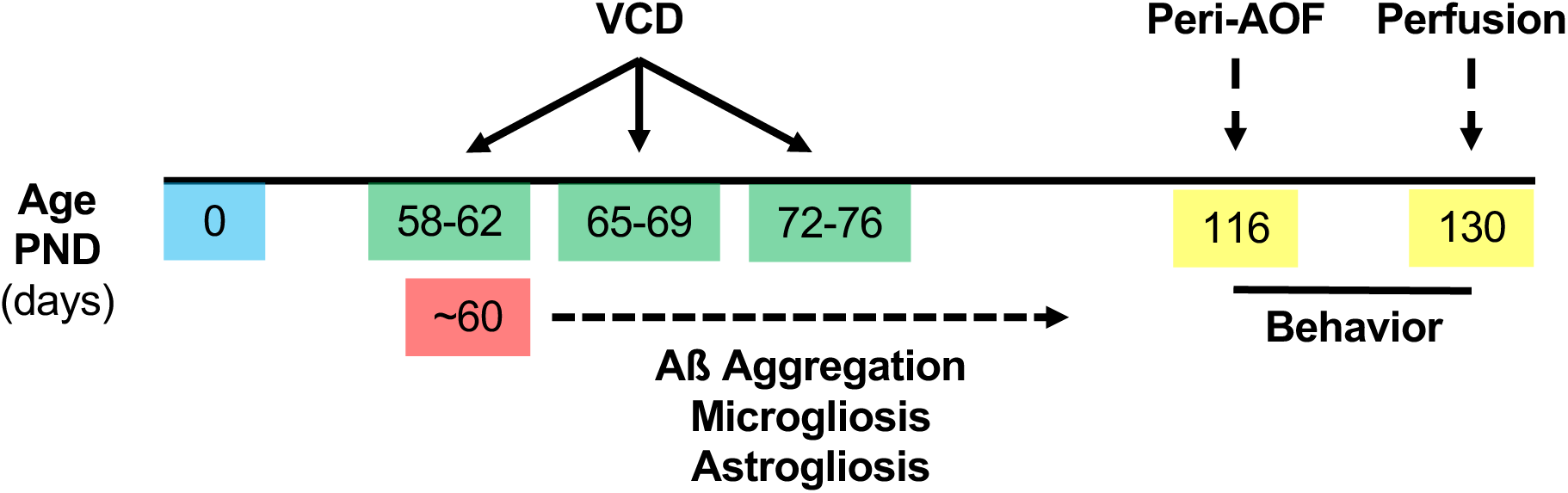
Timeline of experimental procedures. VCD (130 mg/kg, i.p.) was injected for 3 weeks, 5 days per week beginning at approximately postnatal day (PND) 58. The extracellular deposition of Aß plaques was expected to begin at ∼PND 60, accompanied by microgliosis and astrogliosis. Behavioral assessments were performed for 2 weeks following the initiation of the peri-AOF phase. Brains were harvested at ∼PND 130.

##### Estrous cycle assessment

At euthanasia, vaginal smears [53] were collected to determine the terminal estrous cycle stage via cytological examination. Estrous cycle phases were classified as proestrus (high estrogen), estrus (declining estrogen), or diestrus (low estrogen and progesterone). Most females were in estrus or diestrus at euthanasia.

### ANTIBODIES

#### 4G8

A mouse monoclonal antibody raised against amino acid residues 17-24 of beta-amyloid (4G8, Biolegend Cat. # 800701) was employed. This antibody recognizes abnormally processed isoforms as well as precursor APP forms (manufacturer’s instructions), and labels both parenchymal and vascular Aβ aggregates [54–57]

#### GFAP

A rabbit polyclonal antibody (Abcam # ab7260; lot # GR20948-21; RRID:AB_305808) raised against full-length human GFAP was used. On Western blot, this antibody recognized 48 kDa and 55 kDa GFAP bands (manufacturer’s datasheet).

#### Iba1

A rabbit polyclonal antibody raised against a synthetic peptide corresponding to the C-terminus of Iba1 (#SAR6502; 019-19741 FUJIFILM Wako Pure Chemical Corporation) was employed. The antibody reacts with rat, mouse and human Iba1 and recognizes a 17 kDa band protein on Western blot (manufacturer’s datasheet). These antibodies have been used in our prior studies [58, 59].

### BRAIN FIXATION AND HISTOLOGY

Mouse brains were processed for immunocytochemistry using established procedures in our labs [60]. Briefly, mice were deeply anesthetized with sodium pentobarbital (150 mg/kg, i.p.), and then perfused with saline. The brains were extracted, bisected sagittally, and the right hemisphere fixed in 4% paraformaldehyde in 0.1M phosphate buffer (PB, pH 7.4) for 24 hrs on a shaker (70 rpm) at 4°C. The forebrain containing the hippocampus was sectioned (40 µm thick) on a vibratome (VT1000X Leica Microsystems, Buffalo Grove, IL) and stored in cryoprotectant (30% sucrose, 30% ethylene glycol in PB) at -20°C until immunocytochemical processing.

For each experiment, one rostral (-2.00 to -2.70 mm from Bregma [61]) or caudal (-2.90 to -3.50 mm from Bregma [61]) hippocampal section per animal was selected and then punch coded in the cortex. Tissue sections from each experimental group were pooled into single containers to ensure identical reagent exposure [60].

### LIGHT MICROSCOPIC IMMUNOCYTOCHEMISTRY

Hippocampal sections from each genotype/sex group were processed for 4G8, Iba1 or GFAP (n= 11/group). Sections were rinsed in 0.1M Tris-saline (TS; pH 7.6), blocked with 0.5% bovine serum albumin (BSA) in TS for 30 min, and incubated in primary antibodies mouse anti-4G8 (1:4000), rabbit anti-GFAP (1:6000) or rabbit anti-Iba-1 (1:4000) diluted in 0.1% Triton-X and 0.1% BSA in TS for 24-hrs at room temperature followed by 24-hrs at +4°C. Next, sections were rinsed in TS and incubated in either biotin-conjugated goat anti-rabbit IgG (for GFAP and Iba1; #111-065-144, Jackson ImmunoResearch Inc., West Grove, PA; RRID:AB_2337965) or goat anti-mouse IgG (for 4G8; # 115-065-166, Jackson ImmunoResearch Inc.; RRID:AB_2338569) in 0.1% BSA and TS. Sections were washed in TS and incubated in Avidin Biotin Complex (ABC; Vectastain Elite kit, Vector Laboratories, Burlingame, CA) at half the manufacturer’s recommended dilution for 30-min. After rinsing in TS, the bound peroxidase was visualized by reaction in 3,3’-diaminobenzidine (Sigma-Aldrich, St. Louis, MO) and 0.003% hydrogen peroxide in TS for 6-minutes (4G8), 3-min (GFAP), or 8-min (Iba1). All primary and secondary antibody incubations were carried out at 145 rpm, whereas rinses were at 90 rpm on a rotator shaker. Sections were mounted in 0.05 M PB onto gelatin-coated glass slides, dehydrated through an ascending series of alcohol to xylene, and coverslipped with DPX (Sigma-Aldrich).

### IMAGE ACQUISITION AND FIELD DENSITOMETRY

Quantification for 4G8, Iba1 and GFAP labeling in the hippocampus were performed using previously established densitometric methods [62–64]. To ensure unbiased data quantification, the analysis was performed by investigators blinded to experimental conditions. Images were acquired using a Nikon Eclipse 80i microscope with a Micropublisher 5.0 digital camera (Q-imaging, BC, Canada) and IP Lab software (Scanalytics IPLab, RRID: SCR_002775). ImageJ64 software (Image J, RRID:SCR_003070) was used to measure the pixel density within regions of interest (ROI) in defined hippocampal subregions. ROIs within four subregions of the rostral and caudal hippocampus were selected: 1) **CA1**: stratum oriens (SO), pyramidal cell layer (PCL), stratum radiatum (SR) and stratum lacunosum-moleculare (SLM); 2) **CA2/3a**: SO, PCL, near and distal SR; 3) **CA3b**: SO, PCL, stratum lucidum (SLu) and SR; 4) **Dentate gyrus** (**DG)**: the supragranular blade (SG), the infragranular blade (IFG) and the central hilus (Cen) and 5) **Subiculum** (caudal section). Background pixel density from non-labeled regions (e.g., corpus callosum) was subtracted to control for illumination variability and background labeling. Prior studies [62] demonstrated a strong correlation between pixel density and actual transmittance, confirming measurement accuracy.

### BEHAVIORAL ASSESSMENTS

Mice were tested sequentially over two weeks in the Novel Object Recognition, Y maze, and Barnes maze tasks, as described in prior studies [65, 66]. The same investigator conducted all behavioral tests. Testing occurred at the same time each day, with results recorded using ANY-maze (Stoelting Co.). Mice were habituated to the testing room for 2 hours daily for five days before testing. On testing days, mice were acclimated to the room for 1-hour before each session. The order of the Y-maze and Novel Object tests were counterbalanced, with 24-hour rest between the tests. Behavioral apparatuses were cleaned with 70% ethanol between trials.

#### Y-maze

This test was used to assess spatial working memory as previously described [60]. Mice were placed in a three-arm maze (40 cm long, 9.5 cm high, 4 cm wide) diverging at a 120° from the central point, and allowed to explore two arms for 5 min (training). After 30 min, the previously blocked arm was opened, serving as the novel arm (test trial). The sequence of arm entries (spontaneous alternation) and locomotor activity were recorded for 5 minutes. A Spontaneous alternation was defined as entities into all the 3 arms on consecutive occasions, and was manually recorded from the recorded videos. The total number of arms entered during the sessions, which reflect locomotor activity, also was recorded. The maximum alternation was subsequently calculated by measuring the total number of arm entries minus 2 and the percentage of alternation was calculated as ((actual alternation/maximum alternation)×100).

##### Novel Object Recognition (NOR)

The NOR apparatus (height 30cm x width 28cm x length 46cm) consisted of an open field chamber with dim illumination throughout, and it is used to measure spatial and working memory. On day 1 (habituation phase), each mouse was allowed to explore the empty arena for 5-min. On day 2 (familiarization phase), two identical objects (type A) were placed on the floor of the area and the mouse was allowed to explore for 5-min followed by a 30-min rest. For the exploration phase, One of the type A objects was replaced with a novel object (type B) and the mouse was allowed to explore for 5-minutes the familiar and novel objects at the same time. For each phase, the total distance traveled, the average speed, the total object exploration time, and the time spent exploring each one of the two objects was recorded. A discrimination index for day 2 exploration phase was calculated as percentage time spent exploring the novel object out of the total object exploratory time.

##### Barnes maze

The apparatus consisted of a maze with a 10cm cylindrical white start chamber in the middle, multiple hole and one escape hole in the periphery. Mice were trained in the apparatus for four sequential days. Each training day consisted of 4 trials (with 15 min inter-trial intervals) in the following sequence: 1) <u>Adaption period</u>. The mouse was placed in the white start chamber, and a buzzer was switched on for 10-sec. Following, the mouse was guided to an escape hole for 15-20-sec. The buzzer was turned off and the mouse was allowed to stay in the escape box for 2-min. 2) <u>Spatial acquisition period</u>. The mouse was placed in the start chamber and the buzzer was switched on for 10 seconds. After 10 seconds, the start chamber was removed, and the mouse was allowed to move around the maze to find an escape hole (maximum 3-minutes). Immediately after the mouse entered the escape hole, the buzzer was turned off and the mouse was allowed to stay in the tunnel for 1-minute. The average values collected from the spatial acquisition period of all 4 trials in a given training day is used as datapoint in the figures (Fig 8A-D). Twenty-four hours after the last training session, mice underwent the <u>probe trial</u>. For this, the mouse was placed in the maze in the white chamber and the buzzer was switched on. After 10-seconds, the chamber was removed, and the mouse behavior was recorded for 90-seconds. For each mouse, the latency time, errors, total length traveled to find the escape hole were recorded.

### IMAGE ADJUSTMENTS FOR FIGURES

Images were adjusted first for contrast and sharpness in Adobe Photoshop 9.0 (Adobe Photoshop, RRID:SCR_014199). Next, images were imported into Microsoft PowerPoint, where final adjustments to brightness, sharpness and contrast were achieved. Images of different groups in the same figure were similarly adjusted to same brightness and contrast. Adjustments were made to the entire image, none of which significantly altered the appearance of the initial raw image.

### DATA ANALYSIS

Data are presented as means ± SEM. Statistical analyses were conducted using Prism 9 software (Graphpad Prism, RRID:SCR_002798) and significance was set at alpha < 0.05. Group comparisons were performed using analysis of variance (ANOVA; one-, two-, three-way) with Tukey or Sidak’s post-hoc tests. Two-group comparisons used Student’s t-tests. Specific analysis conducted are indicated in figure legends. Graphs were generated in Prism 9 software.

## RESULTS

### Peri-AOF is associated with increased amyloid fibrils in select regions of the CA1 and CA3, but not in the dentate gyrus, in 5xFAD mice

AD dementia is more prevalent in women and may emerge at the onset of menopause. However, there is limited evidence that Aβ levels in the hippocampus, which are a hallmark of AD, are influenced by perimenopause. To evaluate whether Aβ increases in the hippocampus of females at a stage of early ovarian failure, 5xFAD females were treated with VCD, or sesame oil as a control, and Aβ levels quantified at peri-AOF, a stage corresponding to human perimenopause. To obtain a more granular understanding of perimenopause role on AD pathology, analysis was performed across major hippocampal subregions. Given the small size and complex geometric borders of these hippocampal subregions, ELISA and similar methods requiring precise tissue punches were not feasible. Instead, we employed light microscopic immunohistochemistry.

The density of 4G8, a marker of Aβ, was examined in CA1, CA3, DG and subicular subregions in rostral and caudal hippocampal sections of the 5xFAD mice (**Fig. 2A-B**). No immunoreactivity was detected in the WT mice. As described below, 4G8 labeling in the 5xFAD mice varied across hippocampal subregions with sublayer specificity.

**Figure 2:**
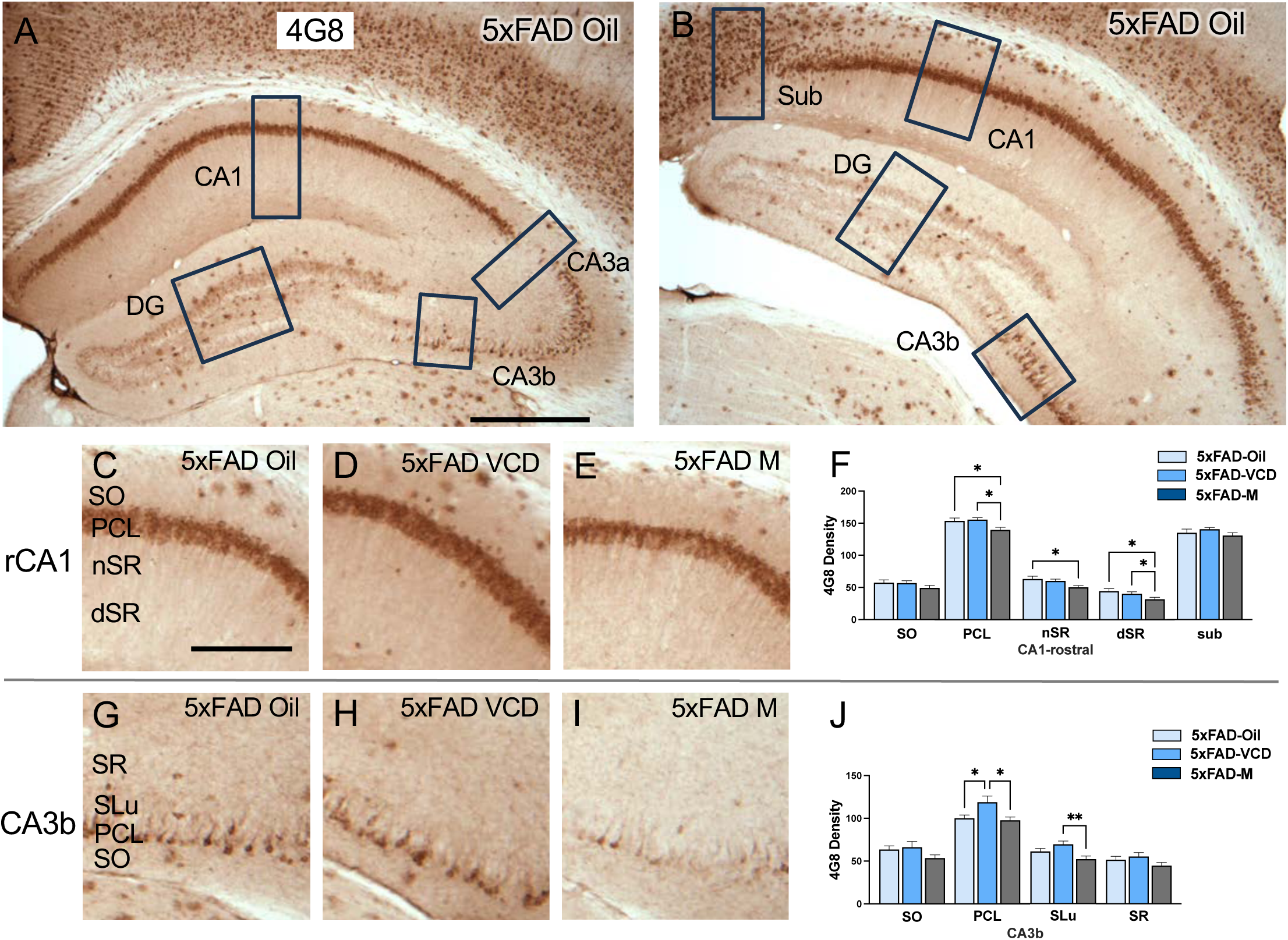
4G8 labeling is differentially altered in select regions of the hippocampus of oil and peri-AOF female and male 5xFAD mice. (A,B) Low-magnification photomicrographs of 4G8 labeling in the rostral **(A)** and caudal **(B)** hippocampus. Boxes indicate regions of the CA1, CA3a, CA3b, dentate gyrus (DG), and subiculum (Sub) that were sampled. **(C,D,E)** Representative photomicrographs showing 4G8 labeling in the rostral CA1 of 5xFAD-oil **(C)**, 5xFAD-VCD **(D)**, and 5xFAD-male mice **(E)**. **(F)** In the rostral CA1 PCL, and dSR, 5xFAD-male mice show significantly less 4G8 labeling than 5xFAD-oil and 5xFAD-VCD female mice. In the rostral CA1 nSR, 5xFAD-male mice show significantly less 4G8 labeling than 5xFAD-oil female mice. **(G,H,I)** Representative photomicrographs showing 4G8 labeling in the CA3b of 5xFAD-oil **(G)**, 5xFAD-VCD **(H)**, and 5xFAD-male mice **(I)**. **(J)** In the CA3b PCL, 5xFAD-VCD female mice show greater 4G8 labeling than both 5xFAD-oil female and 5xFAD-male mice. In the CA3b SLu, 5xFAD-VCD female mice show greater 4G8 labeling than 5xFAD-male mice. * p < 0.05; ** p < 0.01 by One-way ANOVA with Tukey’s post-hoc comparisons. Data are expressed as mean +/-SEM, n = 11 animals per experimental group. Scale bars (A,B) = 500 μm, (C,D,E,G,H,I) = 200 μm.

#### CA1 and subiculum

In the rostral CA1, dense 4G8-immunoreactivity (ir) was found in the pyramidal cell layer (PCL), with scattered clusters in the other laminae, particularly in the stratum oriens (SO) (**Fig. 2C-E**). In the rostral CA1 PCL, 5xFAD male mice exhibited lower 4G8 labeling than both 5xFAD-oil treated and 5xFAD VCD-treated female mice following post-hoc analysis (F=5.080, p=0.013) (**Fig. 2F**). Similarly, in the stratum radiatum (nSR), 5xFAD males showed lower 4G8 labeling compared to 5xFAD-oil treated female mice (F= 3.783, p = 0.034) (**Fig. 2F**). Analogously, in the deep stratum radiatum (dSR), the density of 4G8 labeling following post-hoc analysis was lower in 5xFAD male mice compared to 5xFAD oil-and 5xFAD VCD-treated female mice (F= 4.178, p = 0.025) (**Fig. 2F**). The 4G8 labeling in the caudal CA1 was similar to that seen rostrally. Moreover, scattered 4G8-positive cells were found in the subiculum though no significant effects were found (Suppl. Fig. 1A-D).

#### CA3

Scattered 4G8-labeled cells were found in the PCL of CA3a and CA3b, with labeled clusters dispersed throughout other laminae (**Fig. 2G-I**). No significant difference in 4G8 density was observed in any sublayers of CA3a (Suppl. Fig. 1E-H). However, in the CA3b PCL, 5xFAD VCD females showed greater 4G8 labeling than both 5xFAD oil-treated females and 5xFAD male mice following post hoc analysis (F=5.049, p = 0.013) (**Fig. 2J**). Additionally, in the stratum lucidum (SLu) of CA3b, 5xFAD VCD-treated females showed higher 4G8 density than 5xFAD males (F = 5.98, p = 0.007; t(20) = 3.374, p = 0.003) (**Fig. 2J**).

#### Dentate Gyrus (DG)

In both the rostral and caudal DG, 4G8-labeling appeared in clusters throughout all laminae, especially in the hilus. No significant difference in 4G8 density was observed in any DG sublayers (Suppl. Fig. 1I-P).

### Peri-AOF is associated with increased levels of reactive astrocytes in select regions of the rostral CA1,DG, and caudal CA1 of 5xFAD mice

Astrocytes facilitate the removal of Aβ from the brain parenchyma by mediating efflux into the cerebral vasculature [67]. Their function is influenced by sex, partly through the actions of estrogen signaling via its estrogen receptors [68–70]. Therefore, astrocytes are expected to play an important role in amyloidosis during ovarian failure. However, evidence is limited regarding changes in astrocyte activity across hippocampal subregions in both intact and reproductively compromised females.

To assess astrocyte activation, the density of the astrocytic marker GFAP was examined in CA1, CA3, DG and subicular subregions within rostral and caudal hippocampus (**Fig. 3A-B**) of WT and 5xFAD oil-and VCD-treated female mice, as well as oil-treated male mice. Consistent with our prior studies [58, 59], GFAP-labeled cells were found throughout all lamina of CA1, CA3 and DG, with fewer GFAP-positive cells in the pyramidal and granule cell layers (**Figs. 3,4**). Representative micrographs of GFAP labeling are shown for rostral CA1 (**Fig. 3C-H**), caudal CA1/subiculum (**Fig. 3J-O**), rostral DG (**Fig. 4A-F**), and caudal DG (**Fig. 4H-M**).

**Figure 3:**
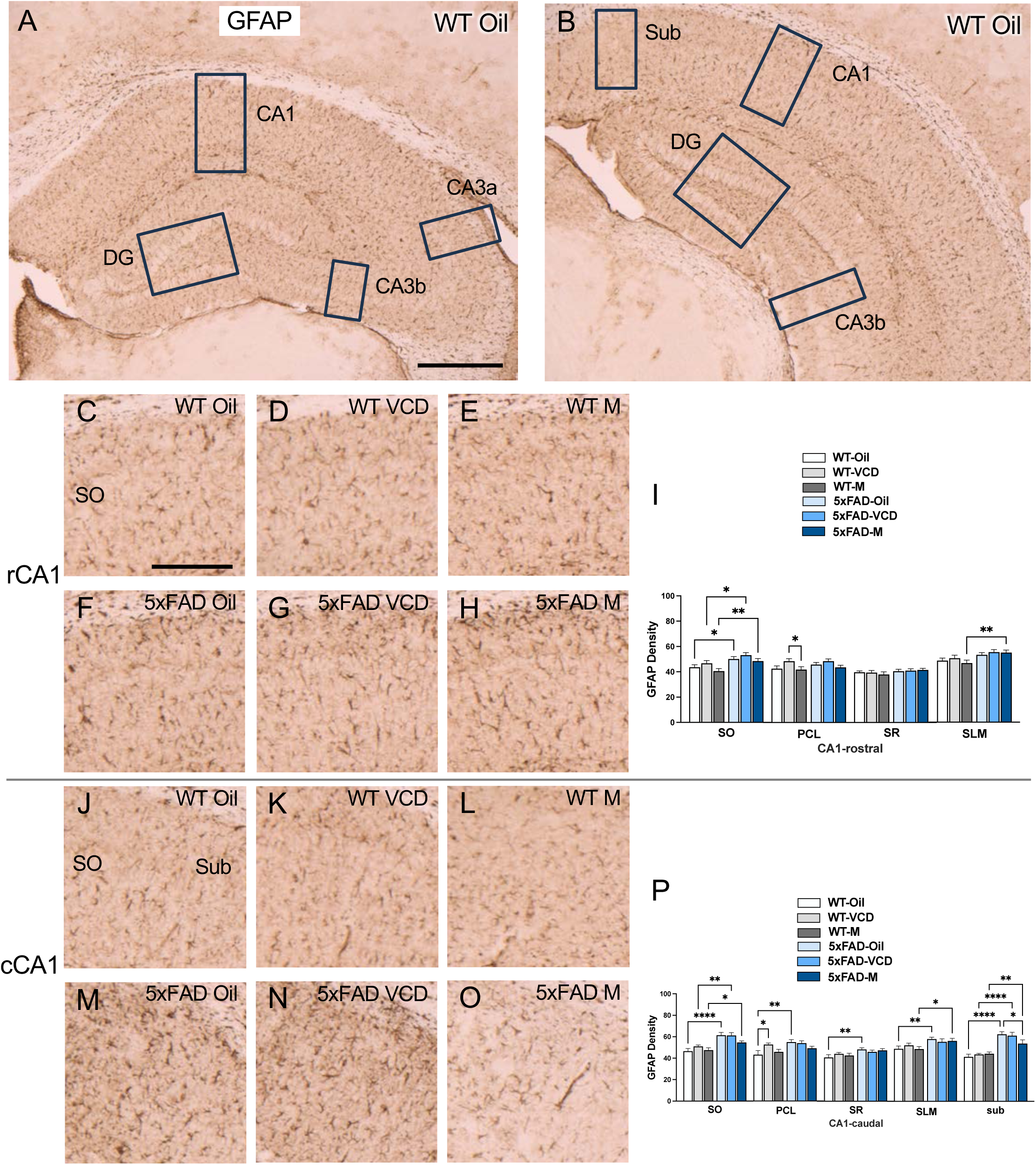
Increased GFAP labeling is associated with peri-AOF and/or 5xFAD genotype in select regions of the hippocampus. **(A,B)** Low-magnification photomicrographs of GFAP labeling in the rostral **(A)** and caudal **(B)** hippocampus. Boxes indicate regions of the CA1, CA3a, CA3b, dentate gyrus (DG), and subiculum (Sub) that were sampled. **(C,D,E,F,G,H)** Representative photomicrographs showing GFAP labeling in the rostral CA1 of WT-oil **(C),** WT-VCD **(D)**, WT-male **(E)**, 5xFAD-oil **(F)**, 5xFAD-VCD **(G)**, and 5xFAD-male mice **(H)**. **(I)** In the rostral CA1 SO, the density of GFAP was increased in 5xFAD mice compared to WT mice, irrespective of sex/AOF treatment. In the PCL, WT-VCD female mice show greater GFAP density than WT-male mice. In the SLM, 5xFAD-male mice show more GFAP labeling than WT-male mice. **(J,K,L,M,N,O)** Representative photomicrographs showing GFAP labeling in the caudal CA1 of WT-oil **(J)**, WT-VCD **(K)**, WT-male **(L)**, 5xFAD-oil **(M)**, 5xFAD-VCD **(N)**, and 5xFAD-male mice **(O)**. **(P)** In the caudal CA1 SO, the density of GFAP was increased in 5xFAD mice compared to WT mice, irrespective of sex/AOF treatment. In the PCL WT-oil female mice show less GFAP labeling than 5xFAD-oil and WT-VCD female mice. In the SR, 5xFAD-oil female mice show greater GFAP density than WT-oil female mice. In the SLM, both 5xFAD-oil female and 5xFAD-male mice had greater GFAP labeling than WT-oil and WT-male mice, respectively. In the Sub, the density of GFAP was increased in 5xFAD mice compared to WT mice, irrespective of sex/AOF treatment, and 5xFAD-oil female mice showed greater GFAP labeling than 5xFAD-male mice. * p < 0.05; ** p < 0.01; **** p < 0.0001 by two-way ANOVA with Tukey’s hoc multiple comparison analysis. Data are expressed as mean +/-SEM, n = 11 mice per experimental group. Scale bars (A,B) = 500 μm, (C,D,E,F,G,H,J,K,L,M,N,O) = 200 μm.

**Figure 4:**
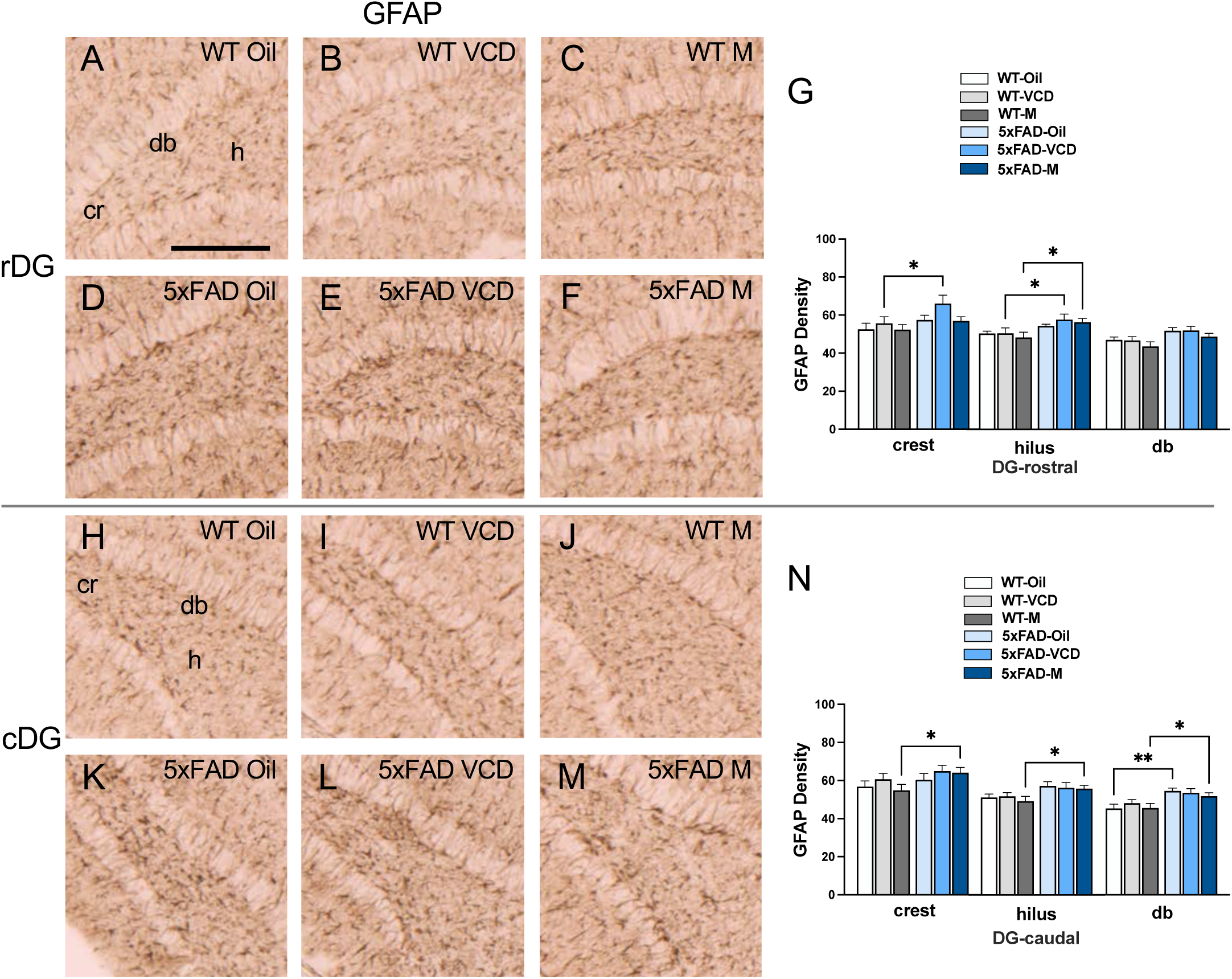
Increased GFAP labeling is associated with peri-AOF and/or 5xFAD genotype in select regions of the hippocampus. **(A,B,C,D,E,F)** Representative photomicrographs showing GFAP labeling in the rostral DG of WT-oil **(A)**, WT-VCD **(B)**, WT-male **(C)**, 5xFAD-oil **(D)**, 5xFAD-VCD **(E)**, and 5xFAD-male mice **(F)**. **(G)** In the rostral DG crest, 5xFAD-VCD female mice show increased GFAP labeling compared to WT-VCD female mice. In the hilus, 5xFAD-VCD female and 5xFAD-male mice show more GFAP labeling than WT-VCD female and WT-male mice, respectively. **(H,I,J,K,L,M**) Representative photomicrographs showing GFAP labeling in the caudal DG of WT-oil **(H)**, WT-VCD **(I)**, WT-male **(J)**, 5xFAD-oil **(K)**, 5xFAD-VCD **(L)**, and 5xFAD-male mice **(M)**. **(N)** In the caudal DG crest, hilus, and db, the density of GFAP was increased in 5xFAD-male mice compared to WT-male mice. In the db, GFAP density was also higher in 5xFAD-oil female mice compared to WT-oil female mice. * p < 0.05; ** p < 0.01 by two-way ANOVA with Tukey’s post hoc multiple comparison analysis. Data are expressed as mean +/-SEM, n = 11 animals per experimental group. Scale bar = 200 μm.

#### CA1 and subiculum

In the rostral CA1 SO region, there was a significant main effect of genotype (F_genotype_ = 18.50, p < 0.0001) and treatment (F_treatment_ = 3.770, p = 0.029). Post-hoc analysis showed that the density of GFAP labeling in SO was greater (p < 0.05) in 5xFAD mice than their WT counterparts (**Fig. 3I**). In the PCL region, a significant main effect of treatment was found (F_treatment_ = 4.822, p = 0.011),with WT-VCD mice showing greater GFAP labeling than WT male mice (p = 0.048) (**Fig. 3I**). In the SLM region, a significant main effect of genotype was observed (F_genotype_ = 12.20, p = 0.0009), and post-hoc analysis showed that 5xFAD male mice had significantly greater GFAP labeling than WT male mice (p = 0.006) (**Fig. 3I**).

In the caudal CA1 SO region, there was a significant main effect of genotype (F_genotype_ = 38.26, p < 0.0001). Post-hoc analysis showed increased GFAP labeling in 5xFAD mice compared to WT controls (p < 0.05) (**Fig. 3P**). In the PLC region, a main effect of genotype was also observed (F_genotype_ = 7.458, p = 0.008). Post-hoc multiple comparisons showed that WT-VCD mice had greater GFAP labeling than WT-oil mice (p = 0.022) (**Fig. 3P**). Additionally, 5xFAD-oil mice had significant higher GFAP labeling in the PLC than WT-oil mice (p = 0.001) (**Fig. 3P**). In the SR region, of caudal CA1, a significant main effect of genotype was observed (F_genotype_ = 9.803, p = 0.003), with increased GFAP labeling in 5xFAD-oil compared to WT-oil mice (p = 0.005) (**Fig. 3P**). In the SLM region, there was a significant main effect of genotype (Fgenotype = 12.49, p = 0.001), with 5xFAD-oil mice exhibiting greater GFAP labeling than WT-oil mice (p = 0.0071), and 5xFAD-male showing increased GFAP compared to WT-male mice (p = 0.0243) (**Fig. 3P**). Similar to other regions of caudal CA1, in the subiculum, a significant main effect of genotype was found (F_genotype_ = 61.46, p < 0.0001), with post-hoc comparisons revealing greater GFAP labeling in 5xFAD mice compared to WT controls (p < 0.05) (**Fig. 3P**). Additionally, 5xFAD-oil mice had significantly higher GFAP labeling in the subiculum than 5xFAD-male mice (p = 0.045) (**Fig. 3P**).

#### CA3

No significant effect of treatment or genotype on GFAP density was observed in any CA3 subregions (Suppl. Fig. 2A-N).

#### DG

In the rostral DG crest, there was a significant effect of genotype (F_genotype_ = 6.914, p = 0.011) on the density of GFAP labeling. Post-hoc analysis revealed significantly greater GFAP labeling in 5xFAD-VCD compared to WT-VCD mice (p = 0.021) (**Fig. 4G**). In the hilus, a significant main effect of genotype was observed (F_genotype_ = 12.21, p = 0.001), with post-hoc multiple comparison analysis revealing that 5xFAD-VCD and 5xFAD-male mice had greater GFAP labeling than their WT counterparts (p < 0.05) (**Fig. 4G**).

In the caudal DG crest, there was a significant main effect of genotype (F_genotype_ = 5.325, p = 0.025), with increased GFAP in 5xFAD-male compared to WT-male mice (p = 0.034) (**Fig. 4N**). In the hilus, a main effect of genotype was also found (F_genotype_ = 10.18, p = 0.002), with 5xFAD-male mice exhibiting greater GFAP labeling than WT-male mice (p = 0.036) (**Fig. 4N**). In the db region of the DG, a main effect of genotype was observed (F_genotype_ = 18.47, p < 0.0001), with increased GFAP labeling in 5xFAD-oil compared to WT-oi mice (p = 0.002) (**Fig. 4N**). Similarly to the crest and hilus, 5xFAD-male mice showed significantly more GFAP labeling than WT-male mice (p = 0.031) in the db (**Fig. 4N**).

These patterns of GFAP labeling suggest that peri-AOF is associated with increased astrocyte activity in select hippocampal regions of the rostral and caudal CA1 and DG.

### Increased microglia activation in select regions of the CA1 and DG of 5xFAD mice

Microglia, the brain’s resident macrophage, regulate neuroinflammation and cognitive function [71]. The protein Iba1 is constitutively expressed in microglia and upregulated upon activation [72, 73], a common feature of aging and neurodegenerative disorders [74]. Significantly, microglia activity is influenced by ovarian hormone changes. Ovariectomy increases Iba1 expression in middle-aged female mice [75], while estradiol reduces microglia reactivity in the hippocampus of aged ovariectomized animals [76]. Additionally, ovariectomy elevates macrophage antigen complex-1, another marker of reactive microglia, in the hippocampus of aged mice [77]. In AD mouse models, chronic estrogen deficiency is linked to heightened microglial activation and neurodegeneration [78]. Given these findings, ovarian failure may alter hippocampal Iba1 expression during amyloidosis, however, direct evidence remains limited.

The density of Iba1 was examined in CA1, CA3, DG and subiculum subregions within the rostral and caudal hippocampus (**Fig. 5A-B**). Consistent with our prior studies [58, 59], Iba1-labeled cells were scattered throughout all lamina in the CA1, CA3 and DG; however, the pyramidal and granule cell layers contained less Iba1 labeling (**Figs. 5 & 6**). Representative micrographs show the distribution of Iba1 labeling in the rostral CA1 (**Fig. 5C-H**), caudal CA1/subiculum (**Fig. 5J-O**), rostral DG (**Fig. 6A-F**), and caudal DG (**Fig. 6H-M**).

**Figure 5:**
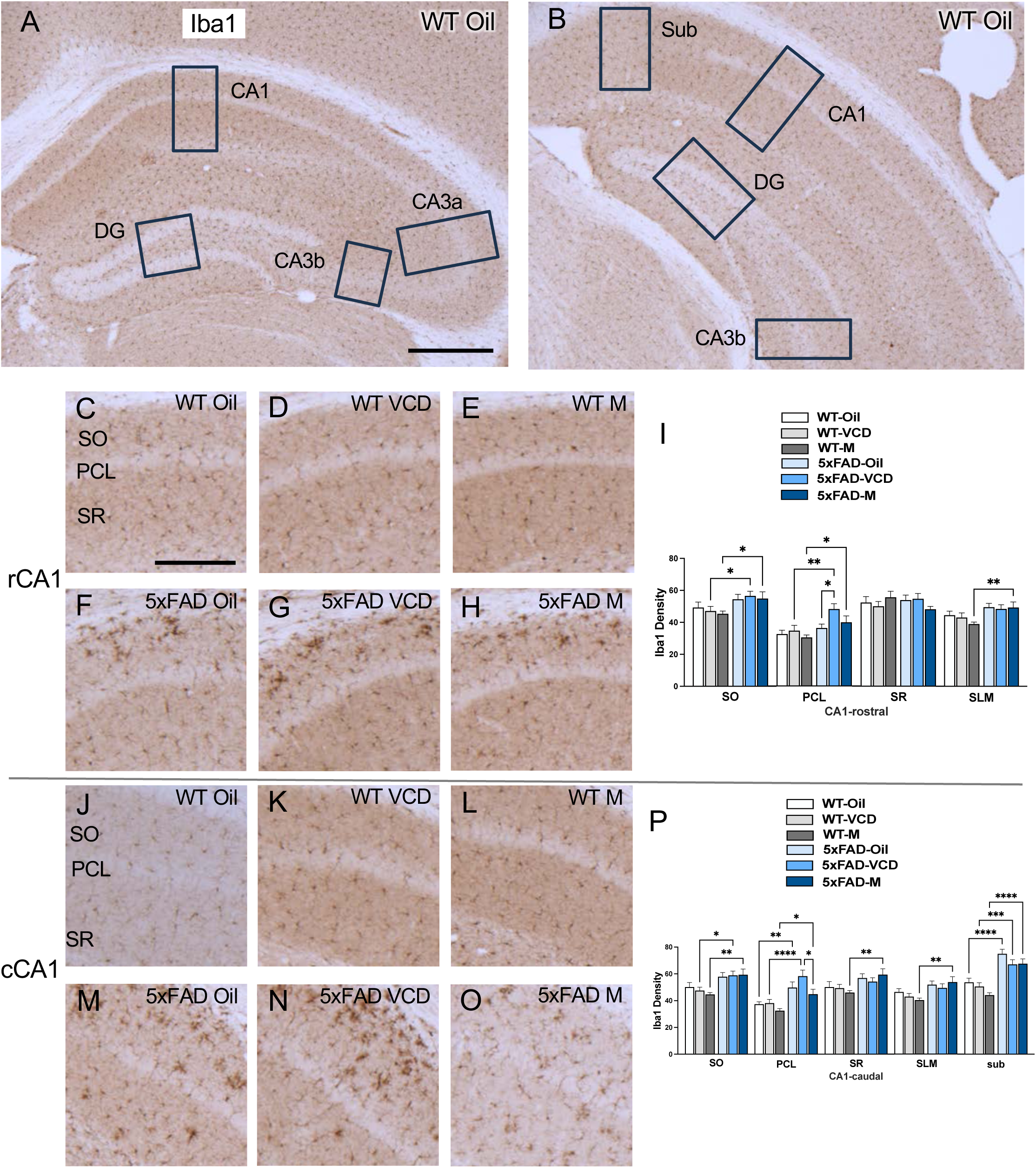
Increased Iba1 labeling is associated with peri-AOF and/or 5xFAD genotype in select regions of the hippocampus. **(A,B)** Low-magnification photomicrographs of Iba1 labeling in the rostral **(A)** and caudal **(B)** hippocampus. Boxes indicate regions of the CA1, CA3a, CA3b, dentate gyrus (DG), and subiculum (Sub) that were sampled. **(C,D,E,F,G,H)** Representative photomicrographs showing Iba1 labeling in the rostral CA1 of WT-oil **(C)**, WT-VCD (D), WT-male **(E)**, 5xFAD-oil **(F)**, 5xFAD-VCD **(G)**, and 5xFAD-male mice **(H)**. **(I)** In the rostral CA1 SO and PCL, the density of Iba1 was increased in 5xFAD-VCD female and 5xFAD-male mice compared to WT-VCD female and WT-male mice, respectively. In the PCL, Iba1 density was greater in 5xFAD-VCD female mice than 5xFAD-oil female mice. In the SLM, 5xFAD-male mice had more Iba1 labeling than WT-male mice. **(J,K,L,M,N,O)** Representative photomicrographs showing Iba1 labeling in the caudal CA1 of WT-oil **(J)**, WT-VCD **(K)**, WT-male **(L)**, 5xFAD-oil **(M)**, 5xFAD-VCD **(N)**, and 5xFAD-male mice **(O)**. **(P)** In the caudal CA1 SO, 5xFAD-VCD female and 5xFAD-male mice had increased Iba1 density compared to WT-VCD and WT-male mice, respectively. In the PCL, the density of Iba1 was increased in 5xFAD mice compared to WT mice irrespective of sex/AOF treatment. 5xFAD-VCD female mice also showed greater Iba1 labeling than 5xFAD-male mice. In the SR and SLM, 5xFAD-male mice had more Iba1 labeling than WT-male mice. In the Sub, 5xFAD mice had increased Iba1 density compared to WT mice irrespective of sex/AOF treatment. * p < 0.05; ** p < 0.01; *** p < 0.001; **** p < 0.0001 by two-way ANOVA with Tukey’s post hoc multiple comparison analysis. Data are expressed as mean +/-SEM, n = 11 animals per experimental group. Scale bars (A,B) = 500 μm, (C,D,E,F,G,H,J,K,L,M,N,O) = 200 μm.

**Figure 6:**
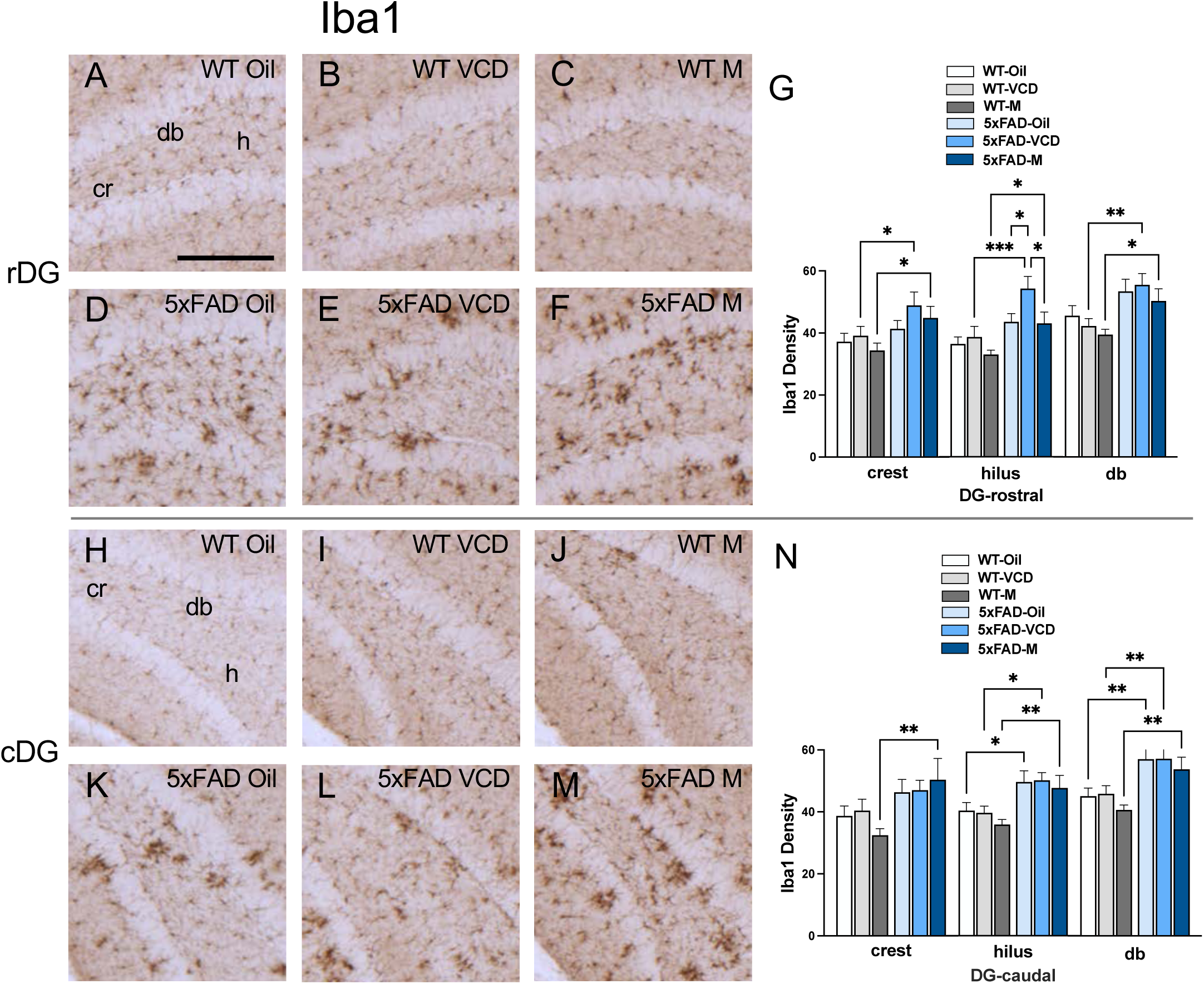
Increased Iba1 labeling is associated with peri-AOF and/or 5xFAD genotype in select regions of the hippocampus. **(A,B,C,D,E,F)** Representative photomicrographs showing Iba1 labeling in the rostral DG of WT-oil **(A)**, WT-VCD **(B)**, WT-male **(C)**, 5xFAD-oil **(D)**, 5xFAD-VCD **(E)**, and 5xFAD-male mice **(F)**. **(G)** In the rostral DG crest and db, 5xFAD-VCD female and 5xFAD-male mice showed greater Iba1 labeling than WT-VCD and WT-male mice, respectively. In the hilus, the density of Iba1 labeling was increased in 5xFAD mice compared to WT mice, irrespective of sex/AOF treatment. 5xFAD-VCD female mice also showed greater Iba1 labeling than 5xFAD-male mice. **(H,I,J,K,L,M)** Representative photomicrographs showing GFAP labeling in the caudal DG of WT-oil **(H)**, WT-VCD **(I)**, WT-male **(J)**, 5xFAD-oil **(K)**, 5xFAD-VCD **(L)**, and 5xFAD-male mice **(M)**. **(N)** In the caudal DG crest, 5xFAD-male mice had greater Iba1 labeling than WT-male mice. In the hilus and db, the density of Iba1 was increased in 5xFAD mice compared to WT mice, irrespective of sex/AOF treatment. * p < 0.05; ** p < 0.01; *** p < 0.001; **** p < 0.0001 by two-way ANOVA with Tukey’s post hoc multiple comparison analysis. Data are expressed as mean +/-SEM, n= 11 animals per experimental group. Scale bar = 200 μm.

#### CA1 and subiculum

In the rostral CA1 SO region, there was a significant main effect of genotype (F_genotype_ = 10.16, p = 0.0023) on Iba1 labeling. Post-hoc analysis showed that Iba1 density was greater (p < 0.05) in 5xFAD-VCD and 5xFAD-male than their WT counterparts (**Fig. 5I**). In the PCL region, there was a main effect of genotype (F_genotype_ = 14.12, p = 0.0004) and treatment (F_treatment_ = 3.449, p = 0.038) on Iba1 labeling. As in the SO region, post-hoc analysis showed increased labeling in 5xFAD-VCD and 5xFAD-male mice (p < 0.05) compared to WT mice (**Fig. 5I**). Additionally, 5xFAD-VCD mice exhibited significantly more Iba1 labeling compared to 5xFAD-oil, mice (p = 0.016) (**Fig. 5I**). In the SLM region of the rostral CA1, a main effect of genotype was observed (F_genotype_ = 10.89, p = 0.002), with post-hoc analysis showing significantly more Iba1 labeling in 5xFAD-male mice than WT-male mice (p = 0.006) (**Fig. 5I**).

In the caudal CA1 SO region, there was a significant main effect of genotype (F_genotype_ = 19.61, p < 0.0001). Post-hoc analysis showed that Iba1 density was greater (p < 0.05) in 5xFAD-VCD and 5xFAD-male mice than their WT counterparts (**Fig. 5P**). In the PCL region, a main effect of genotype (F_genotype_ = 30.99, p < 0.0001) and treatment (F_treatment_ = 4.272, p = 0.014) was observed . Multiple comparisons revealed significantly more Iba1 labeling in 5xFAD mice than in WT mice (p < 0.05) (**Fig. 5P**). Additionally, 5xFAD-VCD mice exhibited greater Iba1 labeling in the PLC than 5xFAD-male mice, (p = 0.014) (**Fig. 5P**). In the SR region of the caudal CA1, a main effect of genotype (F_genotype_ = 9.983, p = 0.003) was found, with post-hoc analysis showing significantly more Iba1 in 5xFAD-male mice than in WT-male mice (**Fig. 5P**). In the SLM, there was also a main effect of genotype (F_genotype_ = 14.19, p = 0.0004), with 5xFAD-male mice exhibiting greater Iba1 labeling than WT-male mice (**Fig. 5P**). Similarly, in the subiculum, a significant main effect of genotype (Fgenotype = 67.47, p < 0.0001) and treatment (F_treatment_ = 3.996, p = 0.024) was found, with post-hoc analysis revealing significantly more Iba1 labeling in 5xFAD mice than WT mice (p < 0.05) (**Fig. 5P**).

#### CA3

There was no effect of treatment or genotype on the density of Iba1 labeling in any of the sublayers of CA3a or CA3b (Suppl. Fig. 2O-AB ).

#### DG

In the rostral DG, there was a significant main effect of genotype (F_genotype_ = 9.923, p = 0.0025) on Iba1 labeling in the crest. Post-hoc analysis showed greater Iba1 density (p < 0.05) in 5xFAD-VCD and 5xFAD-male mice than in WT mice (**Fig. 6G**). In the hilus, there was a main effect of genotype (F_genotype_ = 20.20, p < 0.0001) and treatment (F_treatment_ = 4.349, p = 0.017). Post-hoc analysis showed significantly more Iba1 labeling (p < 0.05) in 5xFAD-VCD and 5xFAD-male mice compared to their WT counterparts (**Fig. 6G**). Additionally, 5xFAD-VCD mice exhibited increased Iba1 density compared to 5xFAD-oil mice (p = 0.036) and 5xFAD-male mice (p = 0.027) (**Fig. 6G**). In the db of the rostral DG, a main effect of genotype (F_genptype_ = 16.39, p = 0.0002) was observed, with post-hoc analysis showing greater density of Iba1 (p < 0.05) in 5xFAD-VCD and 5xFAD-male mice compared to WT mice (**Fig. 6G**).

In the caudal DG, a main effect of genotype (F_genotype_ = 10.04, p = 0.002) was found in the crest, with 5xFAD males exhibiting greater Iba1 density than WT male mice (p = 0.003) (**Fig. 6N**). In the caudal hilus, a significant main effect of genotype (F_genotype_ = 20.42, p < 0.0001) was observed, with post-hoc analysis showing greater Iba1 labeling (p < 0.05) in 5xFAD mice than in their WT counterparts (**Fig. 6N**). Similarly, a significant main effect of genotype (F_genotype_ = 25.07, p < 0.0001) was determined in the db of the DG, with multiple comparisons revealing significantly more Iba1-labeling (p < 0.01) in 5xFAD mice than in WT mice (**Fig. 6N**).

These results suggest that the 5xFAD genotype at peri-AOF influences microglia activation in multiple subregions of rostral and caudal CA1and DG compared to peri-AOF WT mice. Additionally, peri-AOF further exacerbates microglia activation in 5xFAD mice in rostral CA1 PCL and DG hilus, suggesting sex-dependent vulnerability. These findings, in concert with increased GFAP labeling in select hippocampal regions, suggest enhanced susceptibility to neuroinflammation at the intersection of AD and perimenopause.

### Cognitive impairment varies with genotype, sex and AOF

Neuroinflammation is intrinsically linked to the progression of cognitive impairment and dementia [79]. However, the impact of early ovarian failure on cognitive function in VCD-treated 5xFAD mice, and how this compares to males, remains unknown. To address this, we assessed the behavioral consequences of early ovarian decline in 5xFAD mice compared to non-VCD treated females and males using different cognitive tests sensitive to Aβ pathology, including the Y-maze alternation test, novel object recognition, and spatial navigation in the Barnes maze.

#### Y-maze

No significant differences were observed in arm alternation behavior (**Fig. 7A**) or in the total number of arm entries in all 3 arms during the testing period (**Fig. 7B**) between 5xFAD and WT mice. No differences were observed in the % novel arm entries respect to total arm entries and in the time spent in the novel arm (data not shown). This may suggest preservation of short-term working memory or that other cognitive domains might be more affected in 5xFAD mice and in VCD-treated females at a young age.

**Figure 7:**
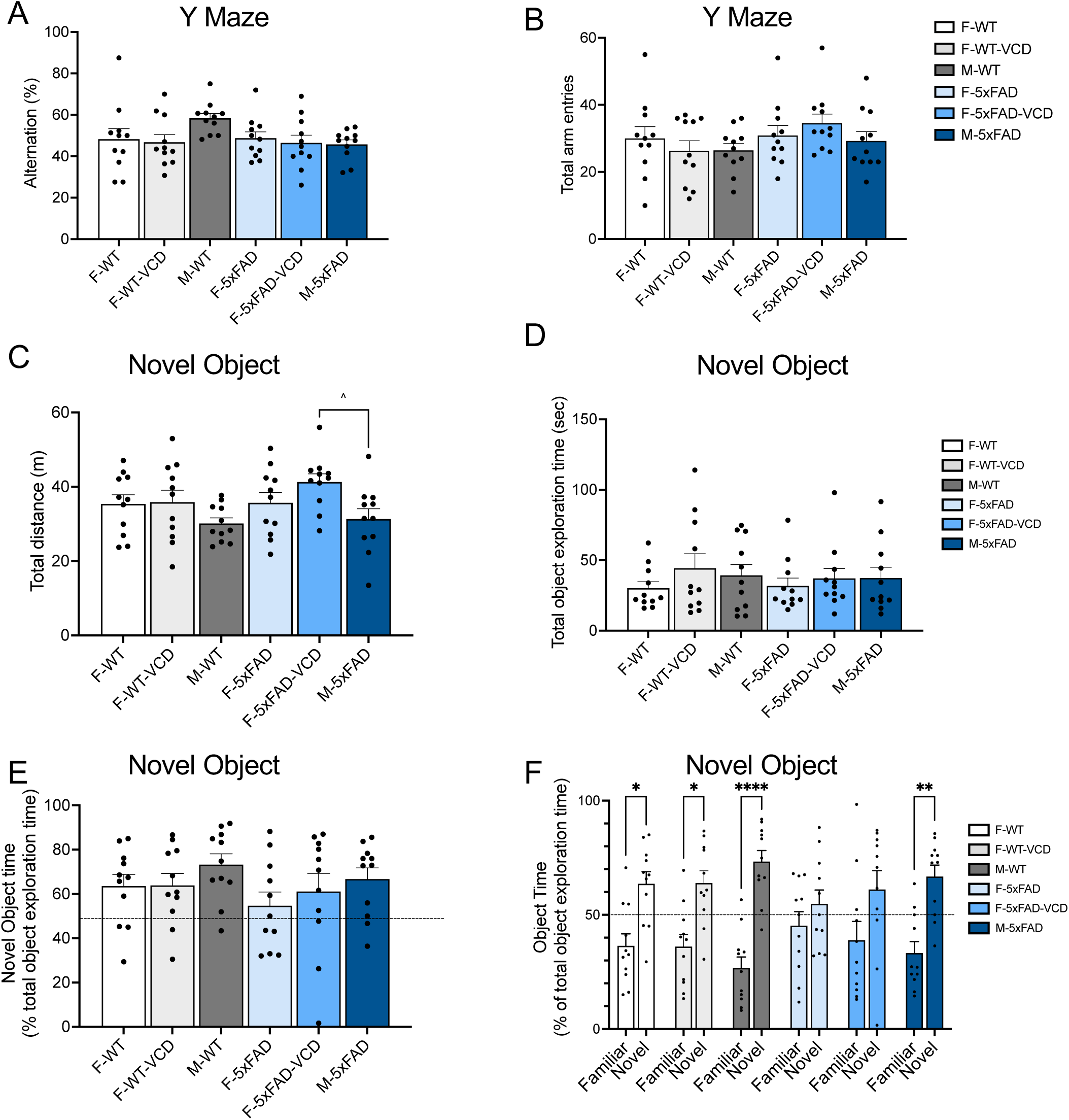
Cognition is not impaired in the Y-maze but is in the novel object recognition test in 5xFAD compared to WT mice. **(A,B)** In the Y-maze test, there were no significant differences in arm alternation behavior **(A)** or in the number of arm entries **(B)** between 5xFAD and WT mice. **(C)** Spontaneous motor activity tended to be decreased in 5xFAD-male mice compared to 5xFAD-VCD female mice. **(D)** total object exploratory time was not changed across experimental groups. **(E)** No significant differences were observed between groups in the percentage time spent exploring the novel object. **(F)** Significant increase in time spent with the novel object vs familiar object was observed for WT male and female (oil and VCD), as well as in 5xFAD male mice. This difference is lost in 5xFAD females. ^ p = 0.06; * p < 0.05; ** p < 0.01; **** p < 0.0001 by one-way ANOVA with Sidak’s post hoc multiple comparison analysis. Data are expressed as mean +/-SEM, n= 11 animals per experimental group.

**Figure 8:**
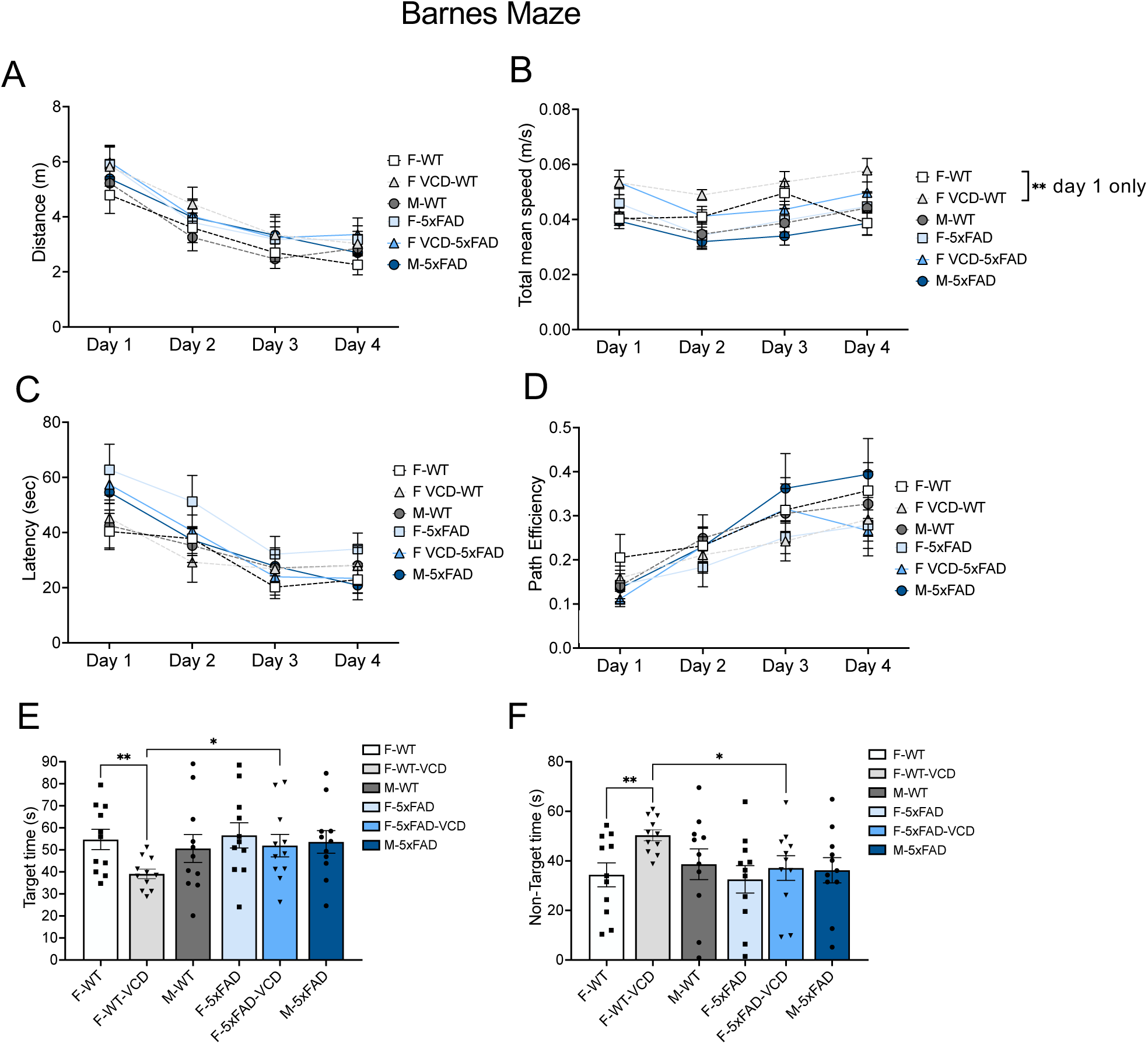
Cognitive performance is differentially affected on the Barnes maze test in peri-AOF female mice. (A,B,C,D) There were no observed differences in motor activity (A,B) and learning performance (C,D) in 5xFAD mice compared to WT mice over time by 2way ANOVA. **(E,F)** WT-VCD female mice demonstrated memory deficits compared to WT-oil and 5xFAD-VCD female mice. * p < 0.05; ** p < 0.01 by 2way ANOVA with Tukey’s post-hoc comparison analysis (A-D) or student’s unpaired t-test (E,F). Data are expressed as mean +/-SEM, *N* = 11 animals per experimental group.

#### Novel object

An effect of AOF was seen in locomotor activity as assessed by total distance travelled (F (5, 60) = 2.42, p = 0.05). 5xFAD-male compared to 5xFAD-VCD female mice had a lower total distance (p = 0.06), although not significant (**Fig. 7C**). No significant differences across groups were observed in the total object exploration time or in Novel Object Recognition Index (**Fig. 7D-E**). Interestingly, significant increase in time spent exploring the novel object vs. the familiar object was observed between WT-male (p < 0.0001) and 5xFAD-male mice (p = 0.001) (**Fig. 7F**). For females, significant differences in time spent exploring familiar and novel objects were found between treatment in WT mice (WT-female (p = 0.018); WT-VCD female (p = 0.014)), but did not extend to 5xFAD females (**Fig. 7F**).

#### Barnes maze

There were no significant differences in motor activity, as assessed by distance traveled and mean speed between 5xFAD and WT mice, regardless of genotype or treatment (**Fig. 8A-B**). Although, a significance difference was observed in mean speed between WT females and VCD-treated WT females at day 1 only (p=0.0086). Also, no differences were observed in learning measures as shown for latency to find the escape and path efficiency (**Fig. 8C-D**) .When cognitive performance was tested 24 hrs after the final acquisition training session, VCD-treated WT-females spent significantly less time in the target quadrant where the escape hole was located, and more time in non-target quadrants compared to both WT-oil or 5xFAD-VCD females (**Fig. 8E-F**). Given that the Barnes maze is a measure of spatial learning and memory, the altered performance in VCD-treated WT females suggests a potential adverse effect of early ovarian failure on spatial memory. This effect appears to be mitigated in 5xFAD-VCD females.

Overall, these behavioral data demonstrate that VCD treatment impairs performance in select cognitive tasks. Combined with our anatomical findings on Aβ deposition and glia activation in the hippocampus, these results suggest that early ovarian failure contributes to early-stage AD-like neurobehavioral pathology.

## DISCUSSION

The relationship between early ovarian failure and neuroinflammation in the hippocampus was investigated using a model of perimenopausal AD that combined chemically-induced AOF with transgenic 5xFAD mice. Age-matched males were also tested in tandem. Mice at approximately 4-months of age showed complex effects on production of Aβ as well as signs of astrocyte and microglia activity that varied by genotype, VCD treatment (AOF) and sex. We found that Aβ expression was elevated in female 5xFAD mice, but only further increased in peri-AOF mice in select hippocampal subregions. Further, increases in glial activation also paralleled Aβ pathology but only in discrete areas of the hippocampus. Assessment of cognitive function showed no effect of peri-AOF in the Y-maze, in the Novel Object Recognition tests, or across Barnes Maze training days (days 1-4). Interestingly, peri-AOF reduced the time spent in the Barnes Maze target quadrant during the testing day (day 5) in WT but not in 5xFAD females, suggesting a treatment-genotype interaction effect. These data provide preclinical experimental evidence supporting the contention that peri-menopause is a sensitive period for neuroinflammation and cognitive function in women.

AD, the most prevalent of the dementias [2], is classically characterized by Aβ deposition and cognitive impairment. However, AD also involves a complex set of neurodegenerative processes, including neuroinflammation [6]. Importantly, the incidence, progression, and severity of the disease are greater in women [9, 10]. Further, perimenopause may be a critical period for the emergence of AD, suggesting that altered gonadal hormone levels contribute to the increased AD risk [11]. Although perimenopause represents a potential turning point in AD pathology, the impact of associated hormonal changes on hippocampal pathology remains unclear. To investigate the role of early ovarian decline in AD pathology, we exposed female 5xFAD mice to the ovotoxin VCD to induce AOF and tested them at an age when brain amyloid deposition and ovarian failure are both at early stages. Age-matched WT mice and 5xFAD male mice were also studied to isolate the impact of AOF independently of amyloidosis and to characterize sex differences, respectively.

We found that Aβ expression was elevated in female 5xFAD peri-AOF mice (injected with VCD) compared to intact females and males injected with sesame oil, in select hippocampal subregions such as the CA1 and CA3b, but not CA3a. The mechanism underlying increased Aß levels in CA3b compared to CA3a in VCD-treated 5xFAD mice is unclear at present. Presumably, these regional differences are mediated by neurophysiological [80, 81], transcriptional [82, 83], connectional [84], morphological [81] and functional [85] divergence across these CA3 subfields, which in turn may be modulated by sex and estrogens. For example, CA3b and CA3a each show unique changes in expression patterns of both stress and plasticity-related molecules following stress, a modulator of dementia [86], and do so in an estrogen-dependent manner [33, 62, 87]. These results suggest that the pathways linking Aß pathology with estrogen signaling within the CA3 are subregion-specific, and further research is needed to clarify their specific contributions to learning and memory during ovarian failure. In VCD-injected WT mice, GFAP density was increased compared to oil-treated WT mice in the caudal CA1 PCL and compared to males in the rostral CA1 PCL. No differences were detected with regard to Iba-1 labeling in any hippocampal region between the WT mice. The increased astrogliosis observed during peri-AOF may result from the loss of astrocyte estrogen signaling. Both resting and activated astrocytes are prominent among the non-neuronal hippocampal cell types that express estrogen receptors alpha and beta [88–90] as well as G-protein estrogen receptors [91]. Further, the expression and cellular location of estrogen receptors in the hippocampus are influenced by estrogen levels [92, 93]. Estrogen modulates GFAP expression [94] and regulates the expression of genes involved in astrocyte proliferation [95]. Importantly, the CA1 PCL is an estrogen receptor-expressing region of the hippocampus [88–90] and is a major hippocampal output that plays a critical role in both spatial and non-spatial memory processes [96]. The functional impact of elevated astrocyte activation in the context of reduced estrogen signaling remains unclear, although it may involve the loss of estrogen-mediated neuroprotection and/or the activation of compensatory mechanisms . This is supported by findings that estrogen reduces glucose- and oxygen-deprivation-induced apoptosis in astrocyte-neuron co-cultures, and that estrogen replacement attenuates the increased apoptosis in the CA1 produced by cerebral ischemia [97]. Given the protective role of estrogen against astrocytosis-associated neural dysfunction, its loss in the CA1 PCL region suggests an increased vulnerability to stress or insult and a heightened propensity for neural pathology.

In addition to their critical metabolic role, astrocytes exert important effects on neuronal signaling and plasticity, particularly via modulation of pyramidal cells [98, 99]. CA1 pyramidal cell hyperexcitability is a hallmark of neurodegeneration and, in particular, is observed in AD models [100–102]. Significantly, astrocytes contribute to CA1 pyramidal cell activity [103–105], and reactive astrocytes are associated with decreased inhibitory synaptic currents and CA1 hyperexcitability [106]. Thus, increased astrocyte activity in the CA1 PCL during early AOF, in the context of disrupted estrogen signaling, may contribute to conditions favoring pyramidal cell dysfunction.

We examined astrogliosis and Aβ expression in peri-AOF 5xFAD mice. Compared to WT VCD-treated mice, 5xFAD VCD mice did not show a further increase in GFAP in the CA1 PCL. However, GFAP density was increased in the CA1 SO and subiculum in VCD-treated 5xFAD mice compared to similarly treated WT mice. Additionally, 5xFAD peri-AOF mice showed higher GFAP density in the crest and hilus of the rostral DG compared to WT mice.

Microglia activation was also observed. Iba-1 levels were higher in peri-AOF 5xFAD mice compared to peri-AOF WT mice in the CA1 PCL, SO and subiculum as well as the different subregions of the DG These results indicate that microglia are recruited in non-pyramidal cell regions of the CA1. Significantly, microglia activation was greater in CA1 PCL and DG hilus cells in 5xFAD peri-AOF mice compared to 5xFAD females and males.

The increased 4G8 density observed in 5xFAD mice was expected to be coupled with elevated GFAP and Iba-1 levels. We found that in VCD-treated mice only the CA3b PCL showed increased Aβ compared to both oil-treated female and male controls. However, there were no differences in either GFAP or Iba-1 across any treatment in any CA3 field. Additionally, in both oil and VCD 5xFAD mice, 4G8 in the CA1 PCL was higher compared to males. Yet, this was associated with increased Iba-1 only in peri-AOF mice, suggesting that this region may be particularly sensitive in response to AOF and Aβ aggregation. The lack of coupling between increased neuroinflammatory markers and Aβ levels in several hippocampal subregions suggests that peri-AOF affects glial activity independently of amyloid aggregation in these areas. Alterations in brain function have been described prior to Aβ plaque or fibril formation in other APP overexpressing mouse models [107]. For example, functional hyperemia is impaired in young mice prior to Aβ aggregation [107] and deficits in autoregulation occur [108]. These effects may be attributed to the actions of soluble oligomeric Aβ [109], which are detectable in 5xFAD mice at the age range studied here [110]. Notably, oligomeric Aβ can impact astrocytes [111] and microglia [112].

Our findings in 5xFAD mice contrast with similar experiments in SwDI mice, a model of cerebral amyloid angiopathy [42, 113]. In SwDI VCD mice, we found increased 4G8 in the CA1 SO and DG. Further, astrocyte and microglia activation was observed in the CA1 SO, and astrocytes in the DG crest. Pyramidal cell layers of the CA1 and CA3 were not affected. These results indicate that dementia models with Aβ pathology in either parenchyma or cerebral vasculature exhibit distinct patterns of hippocampal amyloid deposition and glial cell activation with notable differences in pyramidal and non-pyramidal regions.

The present results demonstrate that distinct hippocampus regions exhibit differential susceptibility to AOF and Aβ in 5xFAD mice. The CA1 of 5xFAD mice, particularly the pyramidal cell layer, was vulnerable to the neuroinflammatory effects of peri-AOF. This aligns with evidence that the CA1 is the earliest hippocampal subregion to exhibit neuronal loss in AD [114, 115] and that astrocyte-pyramidal cell interactions in the CA1 are disrupted in AD models [116].

Interestingly, the pyramidal cell layer of the CA3b region uniquely demonstrated elevated Aβ in 5xFAD peri-AOF mice. The CA3 is a region that expresses estrogen receptors [88, 117–120], contains estrogen-sensitive neurons [33], and may be protected against Aβ-mediated degeneration by estrogen [121]. Together with our findings in WT mice, these results suggest that pyramidal cell layers of the CA1 and CA3 fields are particularly sensitive to decreased estrogen and increased Aβ during early AOF in female mice. In sum, these findings highlight the complex interplay between ovarian hormone loss, neuroinflammation, and amyloid pathology, underscoring the need for further research into the mechanisms by which perimenopause contributes to heightened AD susceptibility in women.

## ABBREVIATIONS

Aβ: beta amyloid
AD: Alzheimer’s disease
AOF: accelerated ovarian failure
cCA1: caudal CA1
Cen: central hilus
cDG: caudal DG
DG: dentate gyrus
GFAP: glial fibrillary acidic protein
Iba1: ionized calcium binding adapter molecule 1
IFG: infragranular blade
PB: phosphate buffer
PCL: pyramidal cell layer
rCA1: rostral CA1
rDG: rostral DG
SG: supragranular blade
SLM: stratum lacunosum-moleculare
Slu: stratum lucidum
SO: stratum oriens
SR: stratum radiatum
TS: Tris saline
VCD: 4-vinylcyclohexene diepoxide

## ACKNOWLEDGEMENTS

Supported by NIH grants R01 HL136520 and HL13650S1 (TAM & MJG), R01 HL135428 (MJG), R01 GM130722 (JP), R21 AG064455 (RM), R01 NS097805 (LP).

## Author contributions

Conceptualization: TAM, MJG, RM, JP, LP; Formal analysis: GS, FY, CW Investigation: GS, FY, TAM, MJG; Resources: TAM, MJG, LP; Writing – original draft preparation: MGJ, TAM, RM, JP; writing - review & editing: GS, RM, JP, MGJ, TAM; visualization: FY, GS; supervision: TAM, MJG; project administration: TAM, MJG; funding acquisition: MJG, TAM, LP.

## FIGURE LEGENDS

**Suppl. Figure 1:**
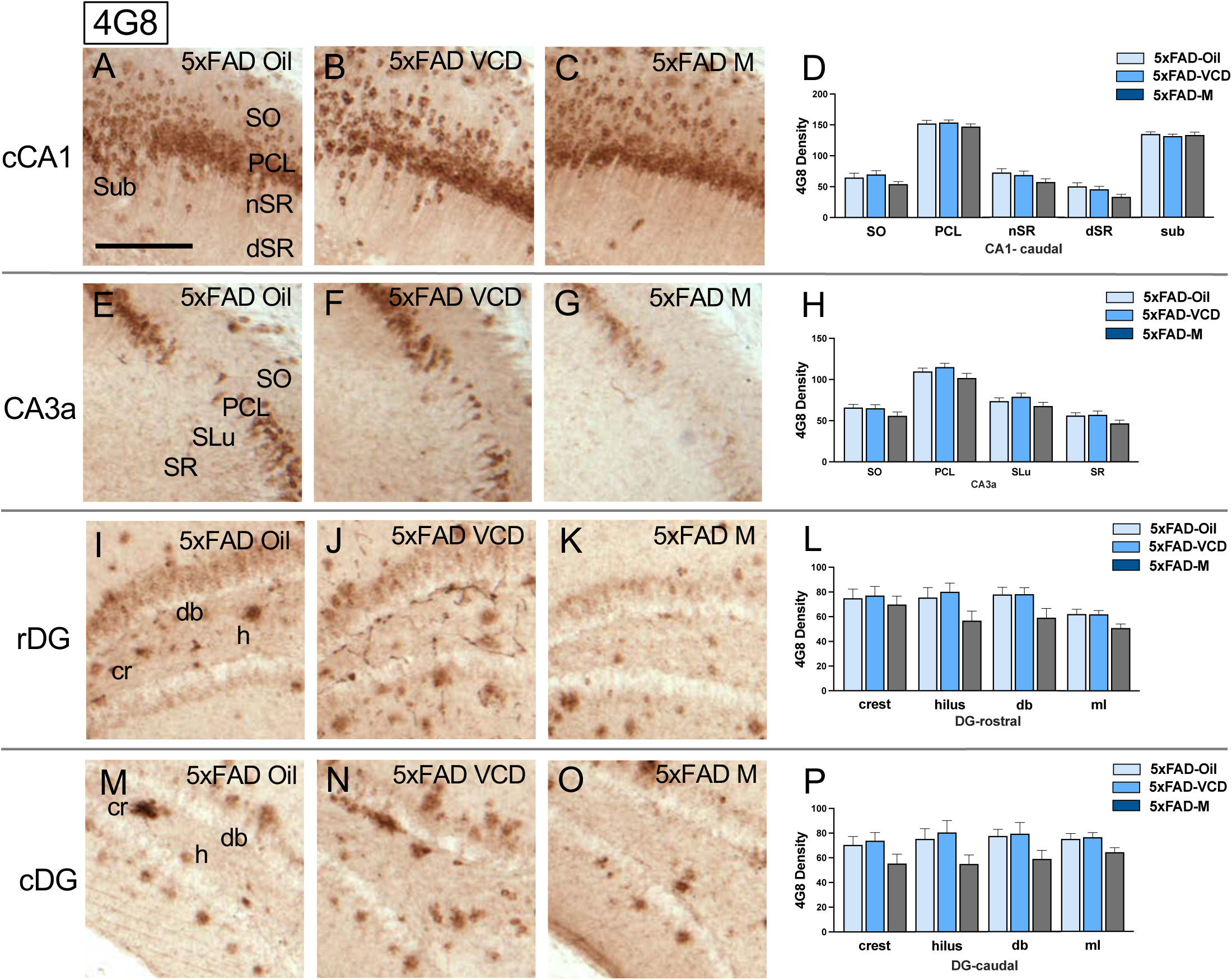
4G8 labeling is not altered in select regions of the hippocampus of across WT and 5xFAD experimental groups. **(A-C)**. Representative photomicrographs showing 4G8 labeling in the caudal CA1 of 5xFAD-oil **(A)**, 5xFAD-VCD **(B)**, and 5xFAD-male mice **(C)**. (**D)** In the caudal CA1, there were no significant differences in 4G8 labeling between 5xFAD-oil, 5xFAD-VCD female, and 5xFAD-male mice. **(E-G)** Representative photomicrographs showing 4G8 labeling in the rostral CA3a of 5xFAD-oil **(E)**, 5xFAD-VCD **(F)**, and 5xFAD-male mice **(G)**. (**H)** In the CA3a, there were no significant differences in 4G8 labeling observed between 5xFAD-oil, 5xFAD-VCD female, and 5xFAD-male mice. (**I-K)** Representative photomicrographs showing 4G8 labeling in the rostral DG of 5xFAD-oil **(I)**, 5xFAD-VCD **(J)**, and 5xFAD-male mice **(K)**. **L.** In the rostral DG, there were no significant differences in 4G8 labeling observed between 5xFAD-oil, 5xFAD-VCD female, 5xFAD-male mice. (**M-O)** Representative photomicrographs showing 4G8 labeling in the caudal DG of 5xFAD-oil **(M)**, 5xFAD-VCD **(N)**, and 5xFAD-male mice **(O)**. (**P)** In the caudal DG, there were no significant differences in 4G8 labeling between 5xFAD-oil, 5xFAD-VCD female, and 5xFAD-male mice. Statistics was calculated by One-way ANOVA with Tukey’s post-hoc multi comparison analysis. Data are expressed as mean +/-SEM, n= 11 mice/group. Scale bar = 200 μm.

**Suppl. Figure 2:**
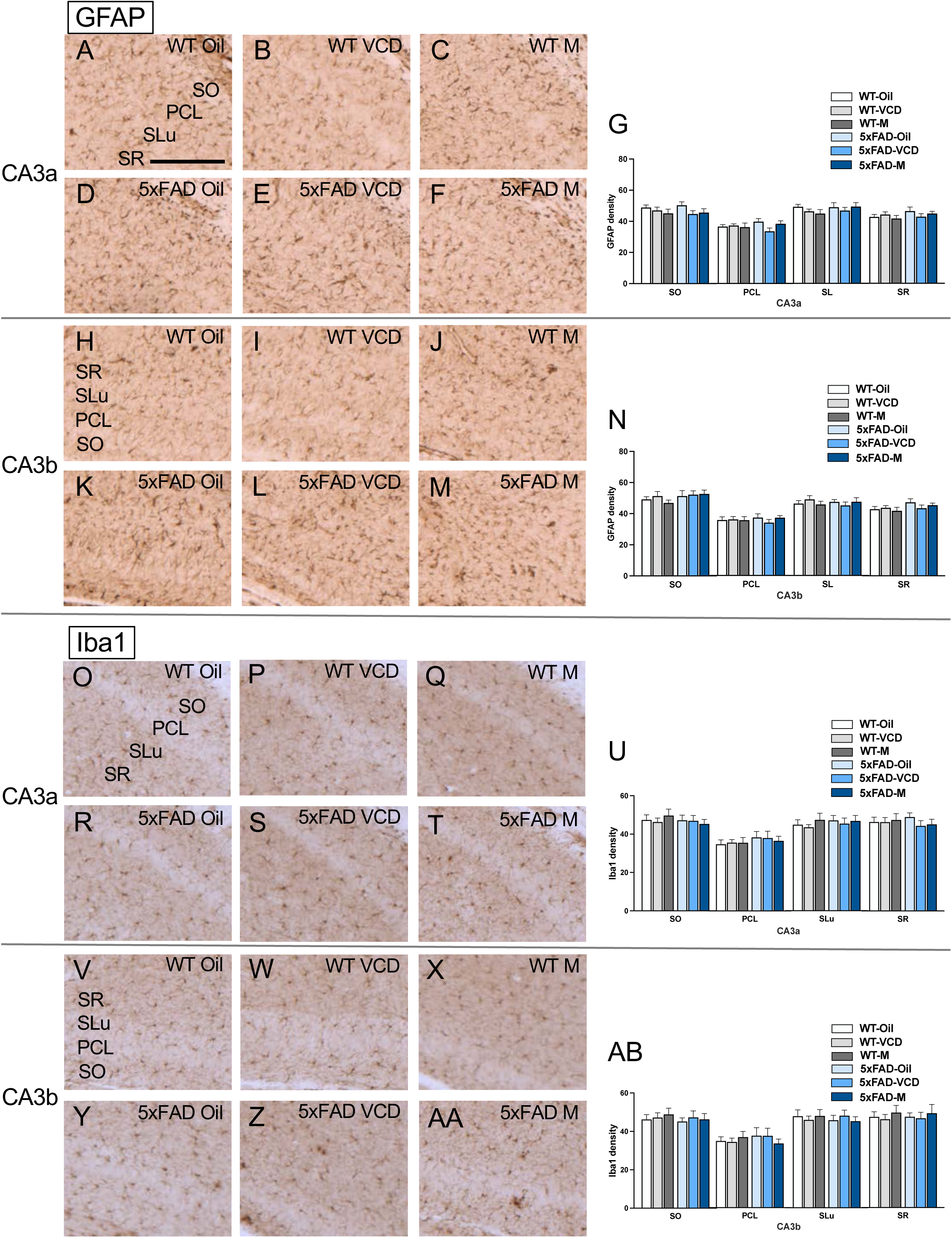
GFAP and Iba1 labeling is unchanged in the CA3a and CA3b regions of the hippocampus of WT and 5xFAD female and male mice. **(A,B,C,D,E,F)** Representative photomicrographs showing GFAP labeling in the rostral CA3a of WT-oil **(A)**, WT-VCD **(B)**, WT-male **(C)**, 5xFAD-oil **(D)**, 5xFAD-VCD **(E)**, and 5xFAD-male mice **(F)**. **(G)** In the CA3a, there were no significant differences in GFAP labeling across groups. H,I,J,K,L,M. Representative photomicrographs showing GFAP labeling in the rostral DG of WT-oil **(H)**, WT-VCD **(I)**, WT-male **(J)**, 5xFAD-oil **(K)**, 5xFAD-VCD **(L)**, and 5xFAD-male mice **(M)**. N. In the CA3b, there were no significant differences in GFAP labeling across groups. **(O,P,Q,R,S,T)** Representative photomicrographs showing Iba1 labeling in the rostral DG of WT-oil **(O)**, WT-VCD **(P)**, WT-male **(Q)**, 5xFAD-oil **(R)**, 5xFAD-VCD **(S)**, and 5xFAD-male mice **(T)**. **(U)** In the CA3a, there were no significant differences in Iba1 labeling across groups. **(V,W,X,Y,Z,AA)**. Representative photomicrographs showing Iba1 labeling in the rostral DG of WT-oil **(V)**, WT-VCD **(W)**, WT-male **(X)**, 5xFAD-oil **(Y)**, 5xFAD-VCD **(Z)**, and 5xFAD-male mice **(AA)**. **(AB)** In the CA3a, there were no significant differences in Iba1 labeling across groups. Statistics was calculated by two-way ANOVA with Tukey’s post hoc multiple comparison analysis. Data are expressed as mean +/-SEM, *n* = 11 animals per experimental group. Scale bar = 200 μm.

